# Identification of novel human topoisomerase III beta inhibitors

**DOI:** 10.1101/2025.03.18.642440

**Authors:** Yasir Mamun, Somaia Haque Chadni, Ramanjaneyulu Rayala, Shomita Ferdous, Rudramani Pokhrel, Adel Nefzi, Prem Chapagain, Yuk-Ching Tse-Dinh

**Affiliations:** Biochemistry PhD Program, Department of Chemistry and Biochemistry, Florida International University; Herbert Wertheim College of Medicine, Center for Translational Science, Florida International University; Department of Physics, Florida International University; Herbert Wertheim College of Medicine, Center for Translational Science, Florida International University; Department of Chemistry and Biochemistry, Florida International University; Biomolecular Sciences Institute, Florida International University; Department of Physics, Florida Interna-tional University; Biomolecular Sciences Institute, Florida International University; Department of Chemistry and Biochemis-try, Florida International University

## Abstract

Human topoisomerase III beta (TOP3B) is a type IA topoisomerase that can change the topology of DNA and RNA substrates via a phosphotyrosine covalent intermediate. TOP3B has been shown to be required for the efficient replication of certain positive-sense ssRNA viruses including Dengue. We applied molecular dynamics simulation combined with docking studies to identify potential inhibitors of TOP3B from a library comprised of drugs that are FDA-approved or undergoing clinical trials for potential drug repurposing. Topoisomerase activity assay of the top virtual screening hits showed that bemcentinib, a compound known to target the AXL receptor tyrosine kinase, can inhibit TOP3B relaxation activity. Additional small molecules that share the *N5*,*N3*-1*H*-1,2,4-triazole-3,5-diamine moiety of bemcentinib were synthesized and tested for inhibition of TOP3B relaxation activity. Five of these molecules showed comparable IC_50_ as bemcentinib for inhibition of TOP3B. However, these five molecules had less selectivity towards TOP3B inhibition versus bemcentinib when inhibition of the type IB human topoisomerase I was com-pared. These results suggest that exploration of tyrosine kinase inhibitors and their analogs may allow the identification of novel topoisomerase inhibitors.

## 1. Introduction

Topoisomerases are essential enzymes required for controlling DNA supercoiling and untangling DNA during vital cellular processes including replication, transcription and repair [1]. Topoisomerase III beta (TOP3B) is the only RNA topoisomerase found in human that can change the topology of both DNA and RNA substrates [2,3] for regulation of genome stability, gene transcription as well as mRNA translation and turnover [4-7]. TOP3B has been shown to be required for efficient replication of many positive-sense single-stranded RNA viruses including flaviviruses (DENV, ZIKV and YFV) [8]. The Dengue virus (DENV) infects up to 400 million people each year as a serious global epidemic with no specific treatment available [9]. The host TOP3B could potentially be hijacked by RNA viruses to assist viral protein translation or viral RNA transport and packaging in different stages of the viral life cycle [10,11]. In addition to use for antiviral therapy, inhibitors of human TOP3B could potentially be useful for anticancer treatment [12]. Many chemotherapeutic agents used in the clinic are topoisomerase poisons that trap the covalent intermediate formed between topoisomerases and chromosomal DNA to induce cancer cell death [13-15]. Specific small molecule inhibitors of type IA topoisomerases that can be used in the clinic remain to be identified. A previous high-throughput screening has identified a bisacridine compound and a thiacyanine compound that can act as TOP3B poisons by acting on RNA to stabilize the TOP3B covalent complex [16]. In this study, we first explored drug repurposing by attempting to identify inhibitors that can target human TOP3B by virtual screening of a library of approved drugs or drug candidates in clinical development. We confirmed experimentally that the tyrosine kinase inhibitor bemcentinib [17,18] can inhibit the catalytic activity of human TOP3B. We then tested additional analogs of bemcentinib for inhibition of TOP3B to compare the potency and selectivity.

## 2. Results

### 2.1. Hits from virtual screening of library of compounds that are FDA-approved or in clinical trials

Molecular docking was performed for 3,855 compounds that are either FDA-approved or in clinical trials against 100 conformations of the TOP3B-RNA covalent complex generated by molecular dynamics simulation. The compounds with the top affinity scores are shown in Table 1. The list reveals that tyrosine kinase inhibitors, with the suffix -tinib, score highly in the computational screening. This is possibly due to the fact that topoisomerases impart their phosphoryl transfer activity through a catalytic tyrosine. These compounds are mostly inhibitors of growth factor receptors (GFR), such as VEGFR-2, and are currently in trial for various cancer treatments [19].

**Table 1.** List of compounds obtained after virtual screening of drugs that are FDA approved or undergoing clinical trial against TOP3B.

| Affinity Score<br>(kcal/mol) | Compound | Known activities<br>(Drugbank description) |
| --- | --- | --- |
| -14.0 | golvatinib | Dual kinase inhibitor of c-Met (hepatocyte growth factor receptor) and VEGFR-2 (vascular endothelial growth factor receptor). |
| -13.8 | bemcentinib | Orally available and selective inhibitor of the AXL receptor tyrosine kinase (UFO), with potential antineoplastic activity. |
| -13.6 | anatibant | Selective and potent Bradykinin B2 receptor antagonist developed for the treatment of traumatic brain injury (TBI). |
| -13.4 | tucatinib | Tyrosine kinase inhibitor that targets the epidermal growth factor receptor 2 (HER2), used in combination therapy against refractory, advanced or metastatic HER2 positive breast and colorectal cancer. |
| -13.4 | phthalocyanine | A dye currently under investigation for treating patients with actinic keratosis, Bowen's disease, skin cancer, or stage I or stage II mycosis fungoides. |
| -13.3 | conivaptan | Arginine vasopressin (AVP) receptor antagonist with affinity for AVP receptor subtypes V1A and V2, approved for hyponatremia (low blood sodium levels) caused by syndrome of inappropriate antidiuretic hormone (SIADH). |
| -13.1 | nilotinib | Tyrosine kinase inhibitor under investigation as a possible treatment for chronic myelogenous leukemia (CML). |
| -13.0 | rebastinib | Orally bioavailable inhibitor of multiple tyrosine kinases with potential antineoplastic activity. |
| -12.9 | radotinib | Second-generation tyrosine kinase inhibitor of Bcr-Abl fusion protein and the platelet-derived growth factor receptor (PDGFR), with potential antineoplastic activity. |

### 2.2. Inhibition of human TOP3B relaxation activity by bemcentinib

From the ranked compounds, we purchased the tyrosine kinase inhibitors golvatinib, bemcentinib, tucatinib, nilotinib, rebastinib and radotinib to test at up to 200 µM concentration for inhibition of the DNA relaxation activity of human TOP3B. Chloroquine was included in the gel electrophoresis buffer to separate the partially relaxed DNA product from the input supercoiled DNA. Inhibition was observed only for bemcentinib (Figure 1 and Figure S1). Figure 1a displays the best pose of bemcentinib docked near the binding pocket of TOP3B. To identify the interactions between bemcentinib and TOP3B, we generated a 2-D interaction map, which shows a polar interaction (3.17Å) between the amide of the 1,2,4-triazole ring of the bemcentinib with the sidechain of Asn519 (Figure 1a, right). We also observed hydrophobic interaction with other nearby residues and the nucleotides towards the 3′ end of the RNA. From serial dilutions, the IC50 of bemcentinib was measured to be 34.3 µM (Figure 1b).

**Figure 1.**
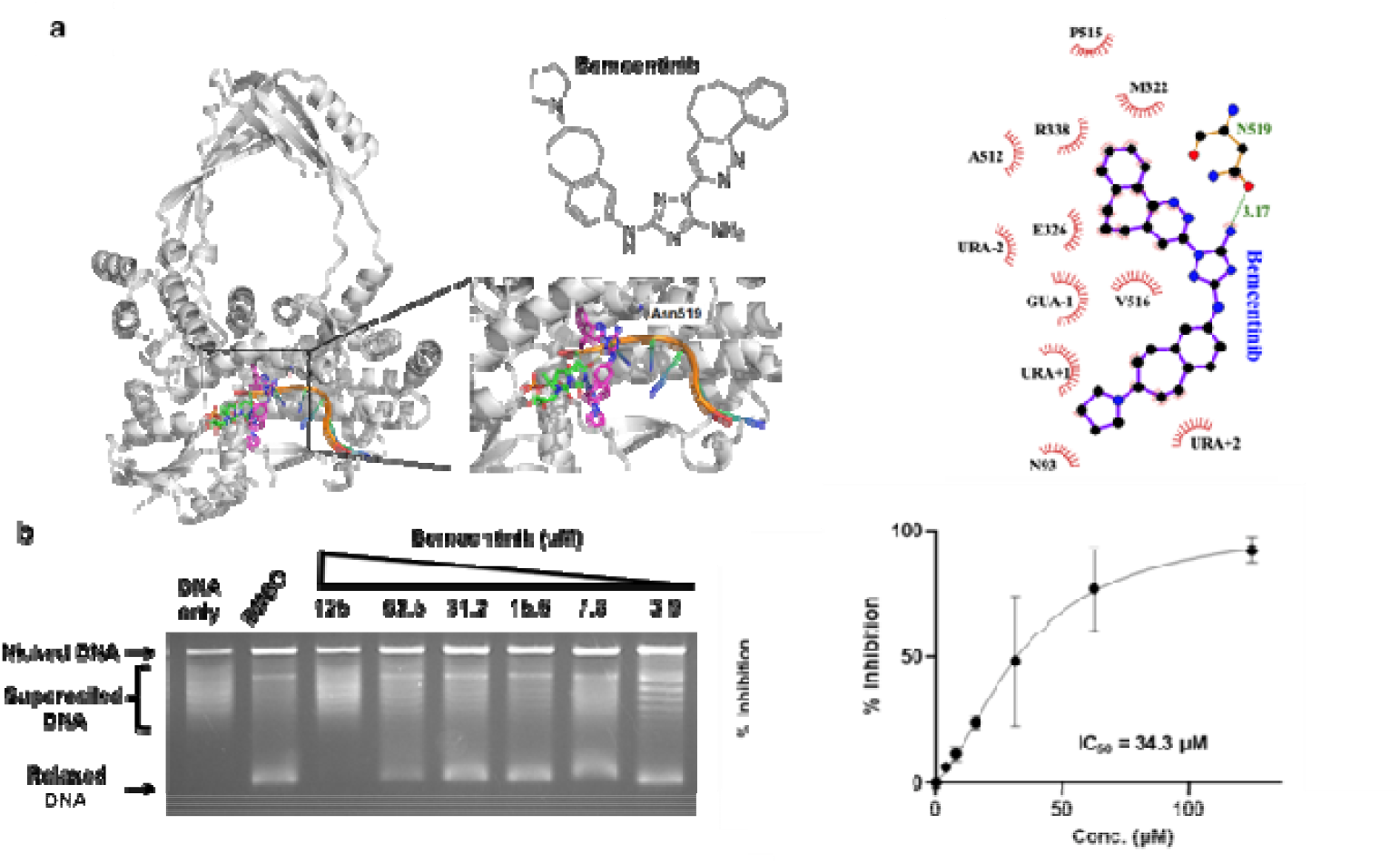
Identification of bemcentinib as a TOP3B inhibitor. (a) The structure of bemcentinib and its docking pose in the target site of TOP3B. Right: Interaction map generated with LigPlot+ for the bemcentinib in its binding site (b) Chloroquine gel electrophoresis assay of TOP3B relaxation activity in the presence of DMSO control or serial dilutions of bemcentinib. The mean and standard deviation of % inhibition from four repeated experiments were plotted with GraphPad Prism version 8.4.2 to obtain the IC_50_ value for bemcentinib. The mean and standard deviation of % inhibition from four repeated experiments were plotted with GraphPad Prism version 8.4.2 to obtain the IC_50_ value for bemcentinib.

### 2.3 Testing of additional small molecules for inhibition of human TOP3B relaxation activity

We synthesized a set of compounds under the 2710 series that have the *N5*,*N3*-1*H*-1,2,4-triazole-3,5-diamine moiety of bemcentinib (Figure 2a). These compounds (Figure S2) were first tested at 200 µM concentration for inhibition of the human TOP3B relaxation activity. Compounds that showed complete inhibition at 200 µM (Figure S3) were tested further at lower concentrations. We found that the compounds shown in Figure 2b could inhibit TOP3B with IC_50_ < 60 µM (Figure 3).

**Figure 2.**
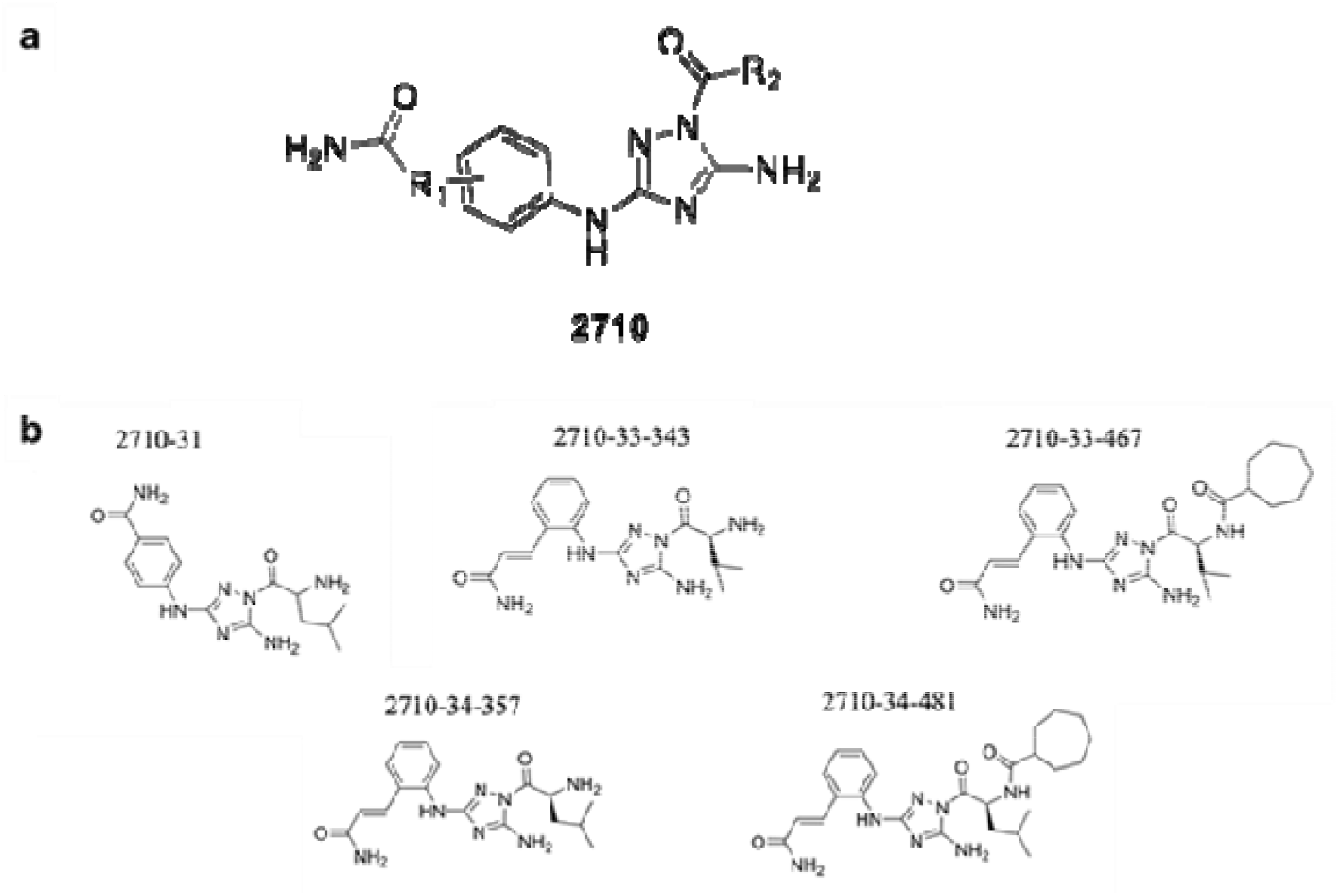
Structure of 2710 series of compounds. (**a**) Scaffold structure with the *N*^*5*^,*N*^*3*^-1*H*-1,2,4-triazole-3,5-diamine moiety. (**b**) 2710-compounds found to inhibit TOP3B with IC_50_ < 60 µM

**Figure 3.**
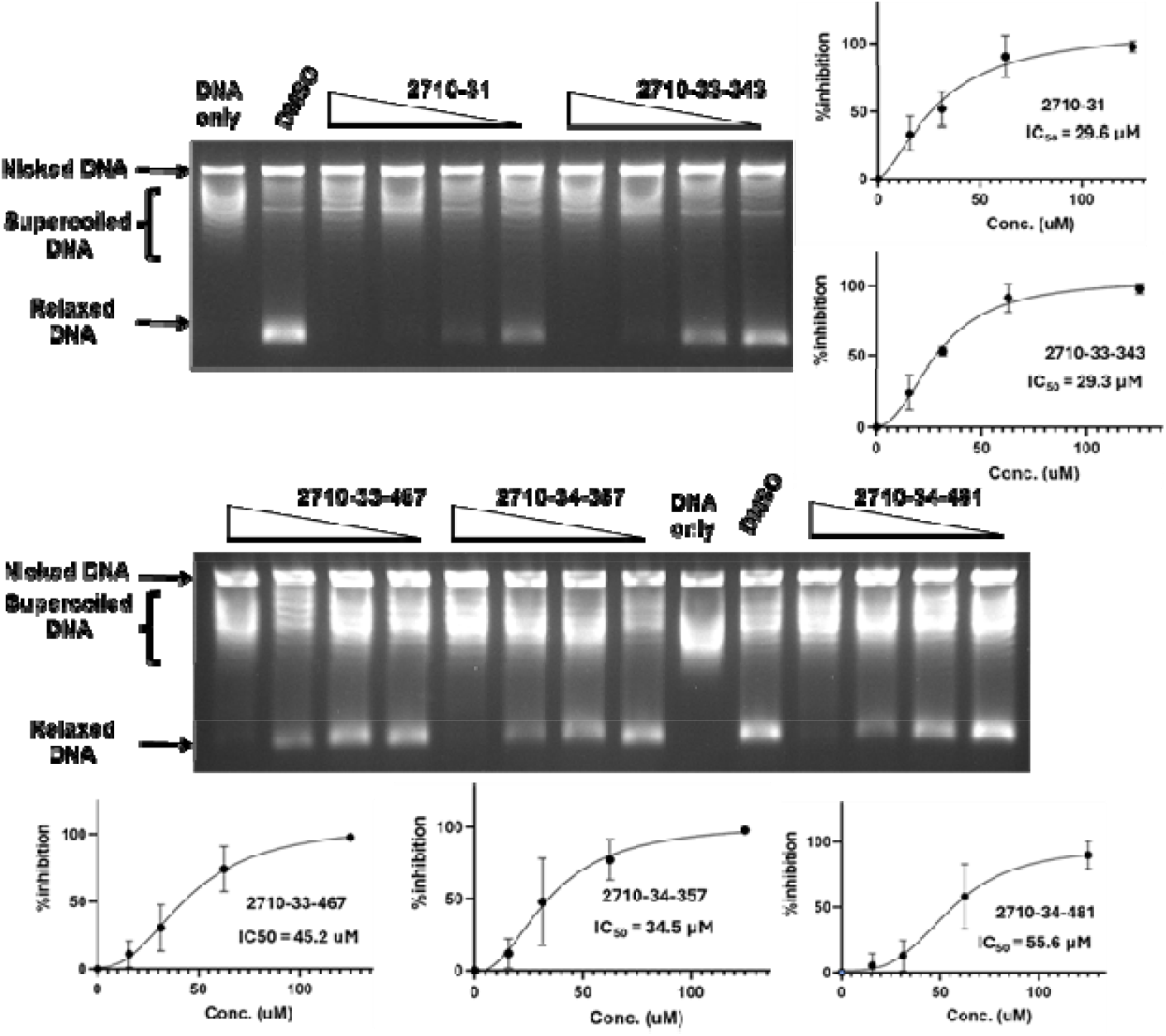
Inhibition of TOP3B relaxation activity by selected compounds from 2710 series. Serial dilution of compounds from 125 to 15.6 µM were assayed by gel electrophoresis in the presence of chloroquine. The mean and standard deviation of % inhibition from three repeated experiments were plotted with GraphPad Prism version 8.4.2 to obtain the IC_50_ values.

### 2.4 Comparison of selectivity for inhibition of human TOP3B versus TOP1

To evaluate the selectivity of bemcentinib and the identified 2710-series compounds as topoisomerase inhibitors, they were also tested for inhibition of type IB human TOP1 which has distinctly different structure and mechanism from the type IA topoisomerase TOP3B. Chloroquine is not required to observe the relaxation of super-coiled DNA by TOP1 by gel electrophoresis because relaxation of supercoiled DNA by TOP1 continues after initial removal of negative supercoils while TOP3B relaxation would not proceed further when the DNA substrate no longer possesses significant single-stranded region. The results of the TOP1 assay (Figure 4, Figure 5) showed that only bemcentinib inhibits TOP3B with greater potency than TOP1 (Table 2). The IC_50_ values for TOP1 inhibition by the 2710-compounds were all lower than the IC_50_ values for TOP3B inhibition.

**Table 2.** IC_50_ values for inhibition of human TOP3B and TOP1 relaxation activity by bemcentinib and inhibitors from the 2710-series.

| Compound | TOP3B IC<sub>50</sub>, μM | TOP1 IC<sub>50</sub>, μM |
| --- | --- | --- |
| bemcentinib | 34.3 | 75.8 |
| 2710-31 | 29.6 | 17.5 |
| 2710-33-343 | 29.3 | 21.3 |
| 2710-33-467 | 45.2 | 29.0 |
| 2710-34-357 | 34.5 | 31.9 |
| 2710-34-481 | 55.6 | 31.1 |

**Figure 4.**
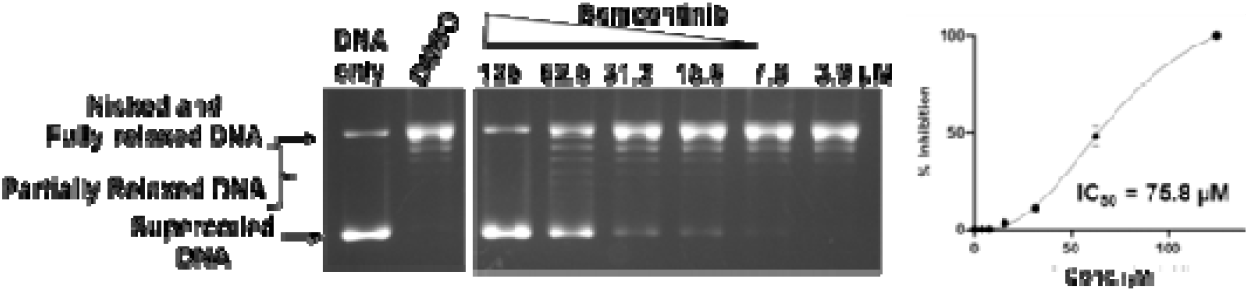
Inhibition of human TOP1 relaxation activity by bemcentinib. The lanes shown are from the same gel. The mean and standard deviation from three repeated experiments were plotted with GraphPad Prism version 8.4.2 to obtain the IC_50_ value.

**Figure 5.**
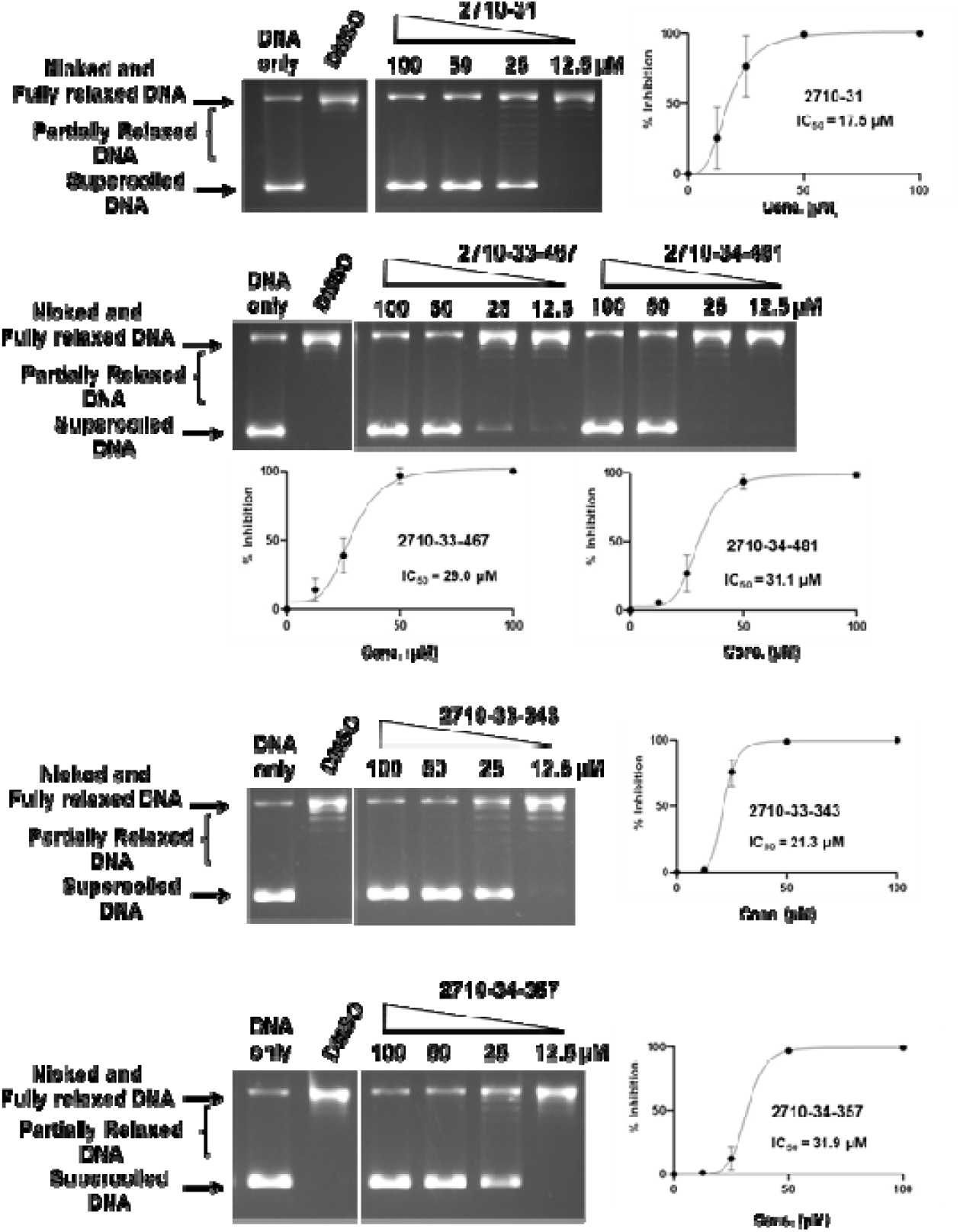
Inhibition of human TOP1 relaxation activity by selected 2710-series compounds. The lanes shown for each compound are from the same gel. The mean and standard deviation of % inhibition from three repeated experiments were plotted with GraphPad Prism version 8.4.2 to obtain the IC_50_ value.

## 3. Discussion

TOP3B is the only human topoisomerase that can change the topology of RNA. Based on a previous report on requirement of human TOP3B for efficient replication of positive-sense ssRNA viruses [8], we conducted virtual screening to try to identify inhibitors of TOP3B from 3,855 approved drugs and drug candidates as potential antiviral candidates. Notably, most of the top virtual screening hits are inhibitors of tyrosine protein kinases. Topoisomerases share similarity with tyrosine protein kinases in likely interactions with a nucleotide moiety as part of ATP for the kinases and part of nucleic acid substrates for topoisomerases, as well as catalysis of Mg(II)-dependent phosphoryl transfer to tyrosine in the reaction mechanism. Dual inhibition of type IA/IIA topoisomerases and tyrosine kinases by small molecules has been reported previously [20,21]. In vitro biochemical assay of six purchasable tyrosine kinase inhibitors identified as TOP3B inhibitor candidates in the virtual screening showed that bemcentinib which targets the AXL receptor tyrosine kinase can inhibit the relaxation activity of TOP3B with

IC_50_ of 34.3 µM. The other five virtual screening hits tested experimentally did not show inhibition in assay at 200 µM. More advanced virtual screening methodology may increase the success rate of future virtual screening projects for TOP3B inhibitors. While it is logical to target the nucleic acid binding sites of topoisomerases for virtual inhibitor screening, these binding sites and interactions involved may not favor specific inhibition by small molecules. Alternative allosteric sites or hinges for conformational change of the topoisomerase could be explored in future virtual screening attempts.

We further explored TOP3B inhibition by the 2710-series of small molecules that possess the *N*^*5*^,*N*^*3*^-1*H*-1,2,4-triazole-3,5-diamine moiety present in bemcentinib. Many of the 2710-series compounds tested showed detectable inhibition of TOP3B relaxation activity at 200 µM, and five of the compounds can inhibit TOP3B with IC_50_ < 60 µM. However, these five compounds were found to also inhibit TOP1 relaxation activity with lower IC_50_. The potency and selectivity for TOP3B inhibition could potentially be further improved by exploring other analogs of bemcentinib. Exploring additional scaffolds of tyrosine kinase inhibitors may also lead to the identification of novel TOP3B or TOP1 inhibitors. More selective TOP3B inhibitors would be useful as tools for basic studies of TOP3B cellular functions or leads for the development of antiviral therapy against flaviviruses that have no currently available treatment options. The potential of the 2710-series compounds identified to be human TOP1 inhibitors could also be further explored for potential discovery of TOP1 poisons for anticancer therapy [13,22] or TOP1 catalytic inhibitors for preventing death from lethal inflammation [23,24].

## 4. Materials and Methods

### 4.1 Molecular dynamics simulation and virtual screening

The TOP3B-RNA covalent complex was generated by first obtaining the apo-structure of the human TOP3B from the protein data bank (PDB ID: 5GVC) [25]. A single stranded RNA with the sequence 5’-AAACUG↓UU-3’ (based on sequence of previously reported TOP3B RNA cleavage oligonucleotide substrate [3]) was placed near the active site of the crystal structure. Next, A covalent bond between the catalytic tyrosine Y336, and the 5’-phosphate of the U nucleotide downstream of the cleavage site in the RNA was computationally generated using a method described previously [26], thus creating a model that resembles the ‘covalently-bound’ structure of the TOP3B enzyme. Conformations were then generated with all-atom molecular dynamics (MD) simulation using standard MD procedure [27] to include possible protein conformational flexibility. The system was prepared using the CHARMM-GUI web interface [28]. The protein was solvated with TIP3 water and 0.15 M NaCl in a cubic box. MD simulation was performed with NAMD 2.13 and NAMD 3.0 [29]. The CHARMM36m [30] force field was used, along with the particle mesh Ewald (PME) [31] method for the long-range ionic interactions and the SHAKE [32] algorithm for constraining hydrogen atom covalent bonds. Nose-Hoover Langevin-piston method with a piston period of 50□fs and a decay of 25□fs was used for pressure control [33], while Langevin temperature coupling with a friction coefficient of 1Lps^−1^ was used for temperature control. The prepared system was minimized for 10,000 steps and equilibrated for 100 ps with 1□fs time step. A 500 ns NPT (constant pressure, temperature) production run was performed at 303.15 K. To prepare the receptor structures for virtual screening, 100 protein conformations were extracted from the production run using VMD [34]. Extracted pdb files were then converted to pdbqt format using AutoDockTools 4.2 [35].

The 3D structures of 3,855 FDA-approved drugs and drugs undergoing clinical trials was obtained in the SDF format from the DrugBank 5.0 database [36]. Then, Open Babel [37] was used to convert the SDF structures to pdbqt format with added polar hydrogen atoms. A cavity that overlaps with the expected nucleic acid binding region near the active site of TOP3B was selected as the target site for the virtual screening, with a box size of 21Å x 25Å x 24Å. Vina from AutoDockTools 4.2 [35] was used to perform docking and screening. Top hits were sorted and ranked based on their binding energy scores using custom scripts. PyMol (The PyMOL Molecular Graphics System, Version 3.0 Schrödinger, LLC) was used for visualization of the best docked pose of bemcentinib. LigPlot+ (v2.2) [38] was used to generate the interaction map between bemcentinib and TOP3B using the best docked pose.

### 4.3 Assay of TOP3B activity inhibition

Golvatinib, bemcentinib, tucatinib, nilotinib, rebastinib and radotinib were purchased from Adooq BioScience. The identity and purity of compounds (>95%) were authenticated by NMR and LC-MS.

Human TOP3B protein was obtained through custom production by Genscript. The coding sequence of human TOP3B was custom synthesized by Genscript and cloned into vector pFastBacGST to generate the recombinant baculovirus expressing human TOP3B protein with N-terminal GST tag and C-terminal 6xHis tag. Following expression in sf9 insect cells, the recombinant TOP3B protein was purified to >95% homogeneity by Genscript using GST column and superdex 200 16/600G chromatography. A buffer containing 20 mM Tris-HCl (pH 7.5), 750 mM KCl, 10% glycerol, 1 mM EDTA, 0.05% NP-40, 1mM DTT was used for storage and dilution.

The presence of TDRD3 protein enhances the relaxation activity of human TOP3B [25,39]. The human TDRD3 coding sequence was amplified by PCR from the TDRD3 clone in pEGFP-C1 [40] using forward primer of 5′-ATGGCCCAGGTGGCCGGC-3′ and reverse primer of 5′-TTAGTTCCGAGCCCGGGGTGG-3′. Amplified TDRD3 gene was joined with the backbone of DHFR-His plasmid (from New England BioLab) using NEB HiFi assembly kit with forward primer: 5′-GGAGGATCCCGGGAATTC-3′ and reverse primer: 5′-GGATCCGTGGTGATGGTG-3′ so that the TDRD3 gene replaced the DHFR gene. The his-tagged TDRD3 protein was expressed from the T7 promoter in *E. coli* Rosetta(DE3) cells and purified using the HisPur Ni-NTA spin column from Thermo Fisher according to the manufacturer’s protocol.

Inhibition of relaxation activity of human TOP3B was carried out in a reaction buffer containing 20 mM HEPES-KOH, pH 7.5, 1 mM DTT, 0.1 mg/ml BSA, and 5 mM MgCl_2_. TOP3B and TDRD3 were added at 18 nM to each reaction mix followed by 0.5 μl DMSO or indicated concentrations of compounds, and lastly by the addition of 500 ng of supercoiled pBAD/Thio plasmid DNA (purified by CsCl gradient centrifugation) for a final reaction volume of 20 μl. After mixing by gentle vortex, the reaction mixtures were spun down and incubated at 37 °C for 1 hour. The relaxation reactions were terminated by the addition of 1 μL of 10% SDS and 2 μL of 800 units/mL proteinase K (New England Biolabs) and incubated at 45□ C for 30 minutes. Following the addition of 5 μL of stop solution (50 mM EDTA, 50% glycerol, and 0.5% v/v bromophenol blue) the reaction products were analyzed in a 1% agarose gel with TAE (40 mM Tris-acetate, pH 8.0, 2 mM EDTA) buffer containing 5 μg/mL chloroquine at 25 V (1 V/cm) for 18 h. Gels were washed with TAE buffer for 1 hour and 50 mM NaCl for 1 hour to remove the chloroquine, and finally with deionized water for 30 minutes. Gels were stained for 2 hours with 1 μg/mL SYBR Gold solution (Thermo Fischer Scientific). Gels were destained with deionized water for 15 min and photographed over UV light with the AlphaImager system. The amount of the relaxed DNA produced by TOP3B in each reaction was quantified using the AlphaImager software version 1.5.0 to calculate the percentage of inhibition versus the DMSO control.

### 4.4 Assay of inhibition of human TOP1 relaxation activity

Recombinant human TOP1 was expressed from clone pYES2-TOP1 transformed into *Saccharomyces cerevisiae strain* EKY3 [41]. Inhibition of TOP1 relaxation activity was assayed as described [41] in a 20 µL reaction containing 200 ng of supercoiled pBAD/Thio plasmid DNA and 1 unit of enzyme in 10 mM Tris-HCl, pH 8.0, 1 mM EDTA, 150 mM NaCl, 0.1 mM spermidine, 0.1 mg/mL BSA and 5% glycerol. The reactions were incubated at 37 °C for 30 min before the addition of 4 µL of stop solution (6% SDS, 0.6% (wv/v)) bromophenol blue and 40% glycerol. Gel electrophoresis for separation of the supercoiled DNA substrate from the relaxed DNA was conducted in 1% agarose gel with TAE buffer for 20 h at 25 V. The DNA was stained with ethidium bromide for an hour and then soaked in water for 5 min. Gels were photographed over UV light with the AlphaImager system. The amount of the supercoiled DNA substrate remaining in each reaction was quantified using the AlphaImager software to calculate the percentage of inhibition versus the DMSO control.

### 4.5 Synthesis of small molecules with disubstituted N^5^,N^3^-1H-1,2,4-triazole-3,5-diamine

The parallel synthesis of all compounds was performed using the previously described “tea-bag” technology [42,43]. The synthetic strategy for the 2710 series of compounds is shown in Scheme 1. Starting from MBHA resin bound nitro aryl compounds, the nitro group was reduced in the presence of tin chloride (SnCl_2_.2H_2_O) and then treated with thiocarbonyldiimidazole (CSIm_2_) in DCM. Following decantation, the generated aryl isothiocyanate was treated with 1*H*-Pyrazole-1-carboxamidine hydrochloride overnight in anhydrous DMF. The resulting thiourea intermediate was treated with hydrazine overnight to generate the corresponding substituted *N*^3^-phenyl-1*H*-1,2,4-triazole-3,5-diamine. Acylation of the obtained compounds with different carboxylic acid led to the desired disubstituted *N*^*5*^,*N*^*3*^-1*H*-1,2,4-triazole-3,5-diamine derivatives. The final compounds were cleaved from the solid support, extracted with acetic acid, and underwent three cycles of freezing and lyophilizing. The crude products were purified by RP-HPLC, and the desired compounds were obtained with purity higher than 85% as determined by LC-MS.

**Scheme 1:**
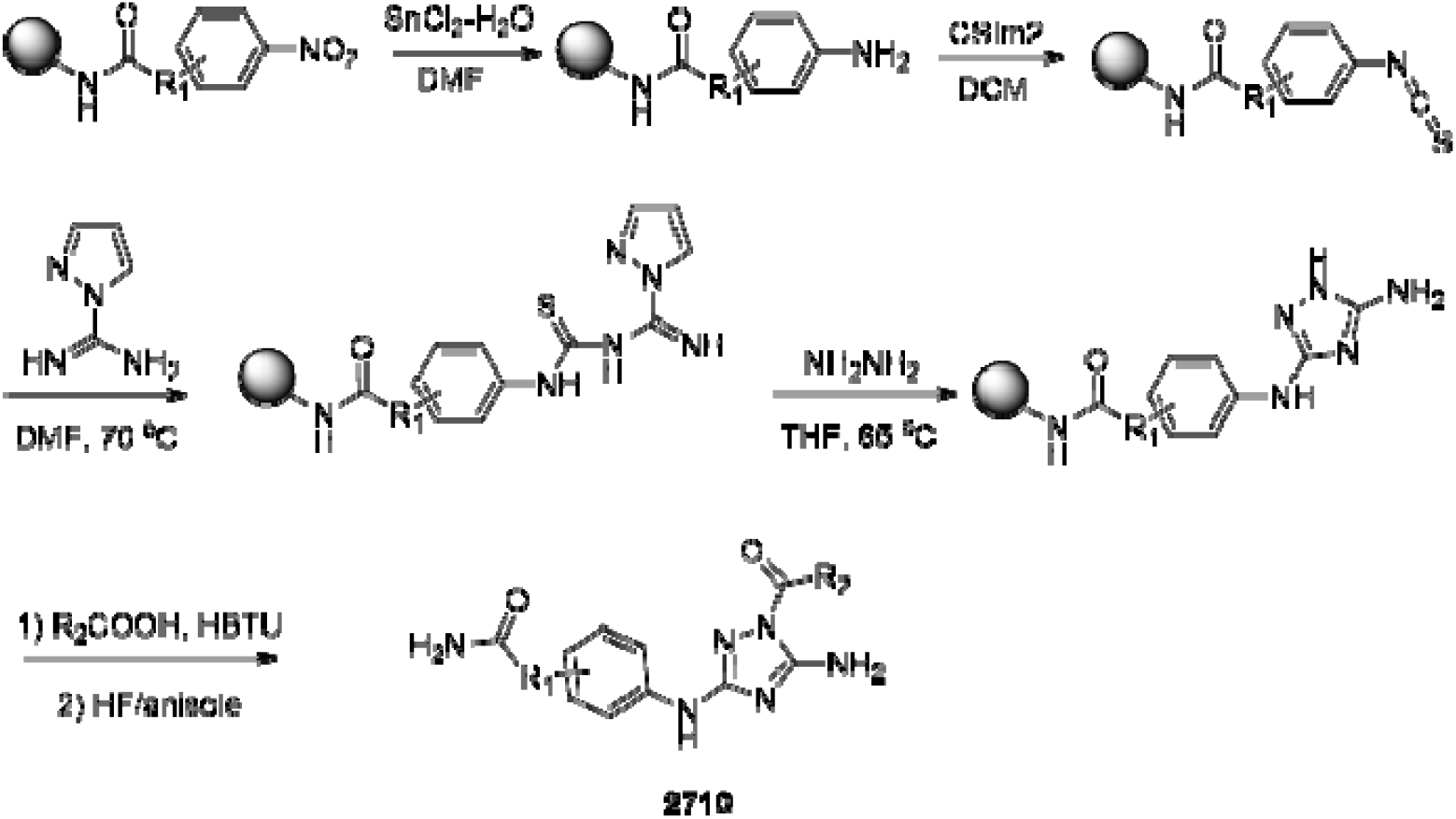
Synthetic strategy of disubstituted *N*^*5*^,*N*^*3*^-1*H*-1,2,4-triazole-3,5-diamine derivatives **2710**.

## 5. Conclusions

In this study, we first identified the tyrosine kinase inhibitor bemcentinib as TOP3B inhibitor from virtual screening of library of approved drugs and drug candidates. After synthesizing and testing the analogs with a similar scaffold, we identified additional TOP3B inhibitors that also inhibit human TOP1 with less selectivity than bemcentinib. The results from this study suggest that investigation of tyrosine kinase inhibitor candidates may yield novel topoisomerase inhibitors.

## Supplementary Materials

Figure S1: Assay of inhibition of human TOP3B relaxation of negatively supercoiled DNA; Figure S2: Structure, chemical formula and molecular weight of 2710 series of compounds tested in this study; Figure S3: Assay of inhibition of human TOP3B relaxation of negatively supercoiled DNA by compounds in the 2710 series; Figure S4: LC/MS of compounds from the 2710 series identified with strongest TOP3B inhibition.

## Author Contributions

Conceptualization, P.C., Y.T. and A.N.; methodology, P.C., A.N. and Y.T.; software, P.C.; formal analysis, Y.M., S.C., R.R., S.F. and R.P.; investigation, Y.M., S.C., S.F. and R.P.; resources, R.R. and A.N.; writing—original draft preparation, Y.M. and Y.T.; writing—review and editing, P.C. and A.N.; visualization, Y.M. and P.C.; supervision, P.C., A.N. and Y.T.; funding acquisition, Y.T. All authors have read and agreed to this version of the manuscript.

## Funding

This research was funded by National Institute of General Medical Sciences of the National Institutes of Health under Award Number R35GM139817 to Y.T. The content is solely the responsibility of the authors and does not necessarily represent the official views of the National Institutes of Health

## Conflicts of Interest

The authors declare no conflicts of interest.

## Notes

### Competing Interest Statement

The authors have declared no competing interest.

## References

1. McKie, S.J.; Neuman, K.C.; Maxwell, A. DNA topoisomerases: Advances in understanding of cellular roles and multi-protein complexes via structure-function analysis. Bioessays 2021, 43, e2000286, doi:10.1002/bies.202000286.

2. Xu, D.; Shen, W.; Guo, R.; Xue, Y.; Peng, W.; Sima, J.; Yang, J.; Sharov, A.; Srikantan, S.; Yang, J.; et al. Top3beta is an RNA topoisomerase that works with fragile X syndrome protein to promote synapse formation. Nat Neurosci 2013, 16, 1238–1247, doi:10.1038/nn.3479.

3. Stoll, G.; Pietilainen, O.P.H.; Linder, B.; Suvisaari, J.; Brosi, C.; Hennah, W.; Leppa, V.; Torniainen, M.; Ripatti, S.; Ala-Mello, S.; et al. Deletion of TOP3beta, a component of FMRP-containing mRNPs, contributes to neurodevelopmental disorders. Nat Neurosci 2013, 16, 1228–1237, doi:10.1038/nn.3484.

4. Zhang, T.; Wallis, M.; Petrovic, V.; Challis, J.; Kalitsis, P.; Hudson, D.F. Loss of TOP3B leads to increased R-loop formation and genome instability. Open Biol 2019, 9, 190222, doi:10.1098/rsob.190222.

5. Su, S.; Xue, Y.; Sharov, A.; Zhang, Y.; Lee, S.K.; Martindale, J.L.; Li, W.; Ku, W.L.; Zhao, K.; De, S.; et al. A dual-activity topoisomerase complex regulates mRNA translation and turnover. Nucleic Acids Res 2022, 50, 7013–7033, doi:10.1093/nar/gkac538.

6. Su, S.; Xue, Y.; Lee, S.K.; Zhang, Y.; Fan, J.; De, S.; Sharov, A.; Wang, W. A dual-activity topoisomerase complex promotes both transcriptional activation and repression in response to starvation. Nucleic Acids Res 2023, 51, 2415–2433, doi:10.1093/nar/gkad086.

7. Tan, K.; Tse-Dinh, Y.C. Variation of Structure and Cellular Functions of Type IA Topoisomerases across the Tree of Life. Cells 2024, 13, doi:10.3390/cells13060553.

8. Prasanth, K.R.; Hirano, M.; Fagg, W.S.; McAnarney, E.T.; Shan, C.; Xie, X.; Hage, A.; Pietzsch, C.A.; Bukreyev, A.; Rajsbaum, R.; et al. Topoisomerase III-ss is required for efficient replication of positive-sense RNA viruses. Antiviral Res 2020, 104874, doi:10.1016/j.antiviral.2020.104874.

9. Sansone, N.M.S.; Marques, L.F.A.; Boschiero, M.N.; Mello, L.S.; Marson, F.A.L. Epidemic after pandemic: Dengue surpasses COVID-19 in number of deaths. Pulmonology 2025, 31, 2448364, doi:10.1080/25310429.2024.2448364.

10. Dasgupta, T.; Ferdous, S.; Tse-Dinh, Y.C. Mechanism of Type IA Topoisomerases. Molecules 2020, 25, doi:10.3390/molecules25204769.

11. Ahmad, M.; Xu, D.; Wang, W. Type IA topoisomerases can be “magicians” for both DNA and RNA in all domains of life. RNA Biol 2017, 14, 854–864, doi:10.1080/15476286.2017.1330741.

12. Zhang, X.; Wang, L.; Zhang, Q.; Lyu, S.; Zhu, D.; Shen, M.; Ke, X.; Qu, Y. Small molecule targeting topoisomerase 3β for cancer therapy. Pharmacol Res 2021, 174, 105927, doi:10.1016/j.phrs.2021.105927.

13. Thomas, A.; Pommier, Y. Targeting Topoisomerase I in the Era of Precision Medicine. Clin Cancer Res 2019, 25, 6581–6589, doi:10.1158/1078-0432.Ccr-19-1089.

14. Bjornsti, M.A.; Kaufmann, S.H. Topoisomerases and cancer chemotherapy: recent advances and unanswered questions. F1000Res 2019, 8, doi:10.12688/f1000research.20201.1.

15. Delgado, J.L.; Hsieh, C.M.; Chan, N.L.; Hiasa, H. Topoisomerases as anticancer targets. Biochem J 2018, 475, 373–398, doi:10.1042/bcj20160583.

16. Wang, W.; Saha, S.; Yang, X.; Pommier, Y.; Huang, S.N. Identification and characterization of topoisomerase III beta poisons. Proc Natl Acad Sci U S A 2023, 120, e2218483120, doi:10.1073/pnas.2218483120.

17. Holland, S.J.; Pan, A.; Franci, C.; Hu, Y.; Chang, B.; Li, W.; Duan, M.; Torneros, A.; Yu, J.; Heckrodt, T.J.; et al. R428, a selective small molecule inhibitor of Axl kinase, blocks tumor spread and prolongs survival in models of metastatic breast cancer. Cancer Res 2010, 70, 1544–1554, doi:10.1158/0008-5472.Can-09-2997.

18. Yadav, M.; Sharma, A.; Patne, K.; Tabasum, S.; Suryavanshi, J.; Rawat, L.; Machaalani, M.; Eid, M.; Singh, R.P.; Choueiri, T.K.; et al. AXL signaling in cancer: from molecular insights to targeted therapies. Signal Transduct Target Ther 2025, 10, 37, doi:10.1038/s41392-024-02121-7.

19. Cohen, P.; Cross, D.; Jänne, P.A. Kinase drug discovery 20 years after imatinib: progress and future directions. Nat Rev Drug Discov 2021, 20, 551–569, doi:10.1038/s41573-021-00195-4.

20. Tse-Dinh, Y.C.; Wong, T.W.; Goldberg, A.R. Virus- and cell-encoded tyrosine protein kinases inactivate DNA topoisomerases in vitro. Nature 1984, 312, 785–786, doi:10.1038/312785a0.

21. Abdelgawad, M.A.; Mohamed, F.E.A.; Lamie, P.F.; Bukhari, S.N.A.; Al-Sanea, M.M.; Musa, A.; Elmowafy, M.; Nayl, A.A.; Karam Farag, A.; Ali, S.M.; et al. Design, synthesis, and biological evaluation of novel pyrido-dipyrimidines as dual topoisomerase II/FLT3 inhibitors in leukemia cells. Bioorg Chem 2022, 122, 105752, doi:10.1016/j.bioorg.2022.105752.

22. Cinelli, M.A. Topoisomerase 1B poisons: Over a half-century of drug leads, clinical candidates, and serendipitous discoveries. Med Res Rev 2019, 39, 1294–1337, doi:10.1002/med.21546.

23. Rialdi, A.; Campisi, L.; Zhao, N.; Lagda, A.C.; Pietzsch, C.; Ho, J.S.Y.; Martinez-Gil, L.; Fenouil, R.; Chen, X.; Edwards, M.; et al. Topoisomerase 1 inhibition suppresses inflammatory genes and protects from death by inflammation. Science 2016, 352, aad7993, doi:10.1126/science.aad7993.

24. Ho, J.S.Y.; Mok, B.W.; Campisi, L.; Jordan, T.; Yildiz, S.; Parameswaran, S.; Wayman, J.A.; Gaudreault, N.N.; Meekins, D.A.; Indran, S.V.; et al. TOP1 inhibition therapy protects against SARS-CoV-2-induced lethal inflammation. Cell 2021, doi:10.1016/j.cell.2021.03.051.

25. Goto-Ito, S.; Yamagata, A.; Takahashi, T.S.; Sato, Y.; Fukai, S. Structural basis of the interaction between Topoisomerase IIIbeta and the TDRD3 auxiliary factor. Sci Rep 2017, 7, 42123, doi:10.1038/srep42123.

26. Tiwari, P.B.; Chapagain, P.P.; Seddek, A.; Annamalai, T.; Uren, A.; Tse-Dinh, Y.C. Covalent Complex of DNA and Bacterial Topoisomerase: Implications in Antibacterial Drug Development. ChemMedChem 2020, 15, 623–631, doi:10.1002/cmdc.201900721.

27. Sandhaus, S.; Chapagain, P.P.; Tse-Dinh, Y.C. Discovery of novel bacterial topoisomerase I inhibitors by use of in silico docking and in vitro assays. Sci Rep 2018, 8, 1437, doi:10.1038/s41598-018-19944-4.

28. Lee, J.; Cheng, X.; Swails, J.M.; Yeom, M.S.; Eastman, P.K.; Lemkul, J.A.; Wei, S.; Buckner, J.; Jeong, J.C.; Qi, Y.; et al. CHARMM-GUI Input Generator for NAMD, GROMACS, AMBER, OpenMM, and CHARMM/OpenMM Simulations Using the CHARMM36 Additive Force Field. J Chem Theory Comput 2016, 12, 405–413, doi:10.1021/acs.jctc.5b00935.

29. Phillips, J.C.; Braun, R.; Wang, W.; Gumbart, J.; Tajkhorshid, E.; Villa, E.; Chipot, C.; Skeel, R.D.; Kalé, L.; Schulten, K. Scalable molecular dynamics with NAMD. J Comput Chem 2005, 26, 1781–1802, doi:10.1002/jcc.20289.

30. Huang, J.; Rauscher, S.; Nawrocki, G.; Ran, T.; Feig, M.; de Groot, B.L.; Grubmüller, H.; MacKerell, A.D., Jr. CHARMM36m: an improved force field for folded and intrinsically disordered proteins. Nat Methods 2017, 14, 71–73, doi:10.1038/nmeth.4067.

31. Essmann, U.; Perera, L.; Berkowitz, M.L.; Darden, T.; Lee, H.; Pedersen, L.G. A smooth particle mesh Ewald method. The Journal of Chemical Physics 1995, 103, 8577–8593, doi:10.1063/1.470117.

32. Ryckaert, J.-P.; Ciccotti, G.; Berendsen, H.J.C. Numerical integration of the cartesian equations of motion of a system with constraints: molecular dynamics of n-alkanes. Journal of Computational Physics 1977, 23, 327–341, doi:10.1016/0021-9991(77)90098-5.

33. Nosé, S.; Klein, M.L. Constant pressure molecular dynamics for molecular systems. Molecular Physics 1983, 50, 1055–1076, doi:10.1080/00268978300102851.

34. Humphrey, W.; Dalke, A.; Schulten, K. VMD: visual molecular dynamics. J Mol Graph 1996, 14, 33–38, 27-38, doi:10.1016/0263-7855(96)00018-5.

35. Trott, O.; Olson, A.J. AutoDock Vina: improving the speed and accuracy of docking with a new scoring function, efficient optimization, and multithreading. J Comput Chem 2010, 31, 455–461, doi:10.1002/jcc.21334.

36. Wishart, D.S.; Feunang, Y.D.; Guo, A.C.; Lo, E.J.; Marcu, A.; Grant, J.R.; Sajed, T.; Johnson, D.; Li, C.; Sayeeda, Z.; et al. DrugBank 5.0: a major update to the DrugBank database for 2018. Nucleic Acids Res 2018, 46, D1074–d1082, doi:10.1093/nar/gkx1037.

37. O’Boyle, N.M.; Banck, M.; James, C.A.; Morley, C.; Vandermeersch, T.; Hutchison, G.R. Open Babel: An open chemical toolbox. J Cheminform 2011, 3, 33, doi:10.1186/1758-2946-3-33.

38. Laskowski, R.A.; Swindells, M.B. LigPlot+: multiple ligand-protein interaction diagrams for drug discovery. J Chem Inf Model 2011, 51, 2778–2786, doi:10.1021/ci200227u.

39. Yang, X.; Saha, S.; Yang, W.; Neuman, K.C.; Pommier, Y. Structural and biochemical basis for DNA and RNA catalysis by human Topoisomerase 3β. Nat Commun 2022, 13, 4656, doi:10.1038/s41467-022-32221-3.

40. Yang, Y.; Lu, Y.; Espejo, A.; Wu, J.; Xu, W.; Liang, S.; Bedford, M.T. TDRD3 is an effector molecule for arginine-methylated histone marks. Mol Cell 2010, 40, 1016–1023, doi:10.1016/j.molcel.2010.11.024.

41. Seddek, A.; Madeira, C.; Annamalai, T.; Mederos, C.; Tiwari, P.B.; Welch, A.Z.; Tse-Dinh, Y.-C. A Yeast-Based Screening System for Differential Identification of Poisons and Suppressors of Human Topoisomerase I. FBL 2022, 27, doi:10.31083/j.fbl2703093.

42. Nefzi, A.; Ostresh, J.M.; Yu, Y.; Houghten, R.A. Combinatorial chemistry: libraries from libraries, the art of the diversity-oriented transformation of resin-bound peptides and chiral polyamides to low molecular weight acyclic and heterocyclic compounds. J Org Chem 2004, 69, 3603–3609, doi:10.1021/jo040114j.

43. Tantak, M.P.; Rayala, R.; Chaudhari, P.; Danta, C.C.; Nefzi, A. Synthesis of Diazacyclic and Triazacyclic Small-Molecule Libraries Using Vicinal Chiral Diamines Generated from Modified Short Peptides and Their Application for Drug Discovery. Pharmaceuticals (Basel) 2024, 17, doi:10.3390/ph17121566.

