## Supplementary Information for "Identification of novel human topoisomerase III beta inhibitors"

**Figure S1. Assay of inhibition of human TOP3B relaxation of negatively supercoiled DNA.** Puchased hits from virtual screening were tested at concentrations of 200, 100 and 50 μM.

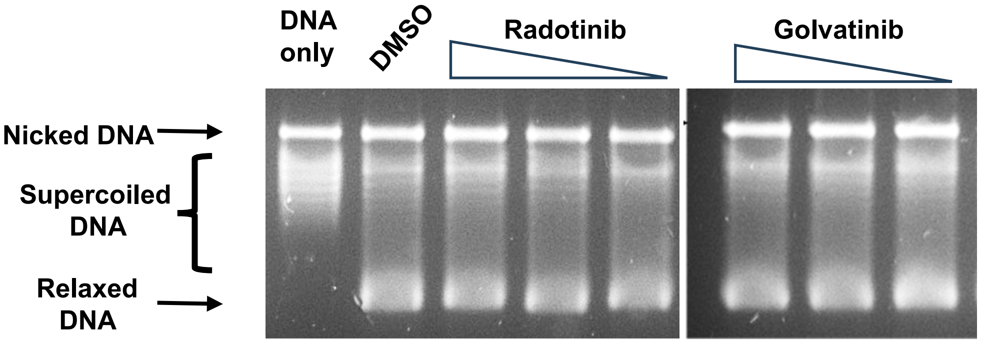


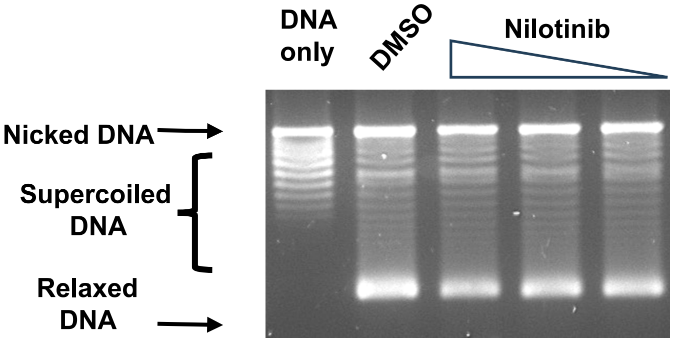


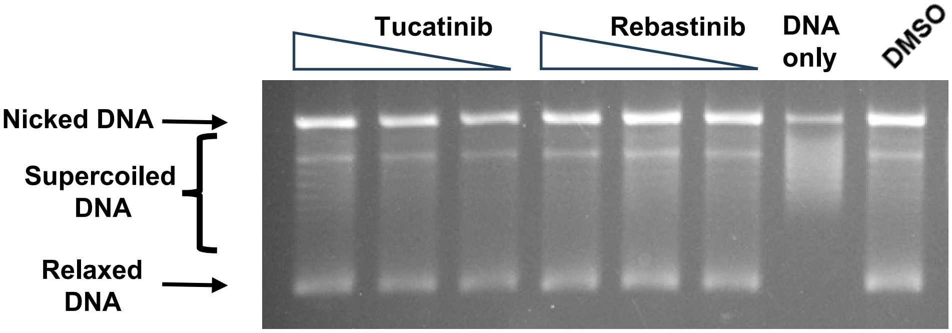

**Figure S2. Structure, chemical formula and molecular weight of 2710 series of compounds tested in this study.**

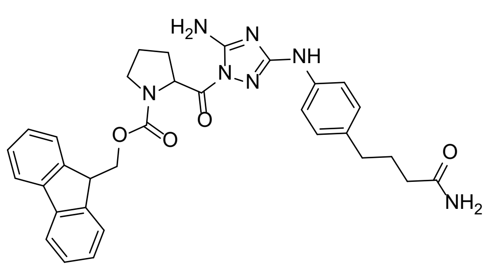

**2710-1**

Chemical Formula: $C_{32}H_{33}N_7O_4$
Molecular Weight: 579.66

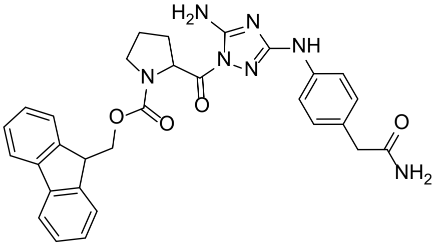

**2710-2**

Chemical Formula: $C_{30}H_{29}N_7O_4$
Molecular Weight: 551.61

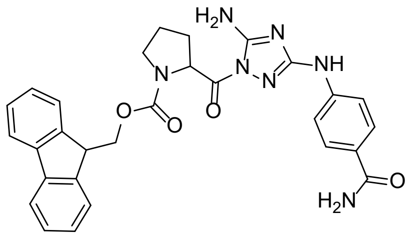

**2710-3**

Chemical Formula: $C_{29}H_{27}N_7O_4$
Molecular Weight: 537.58

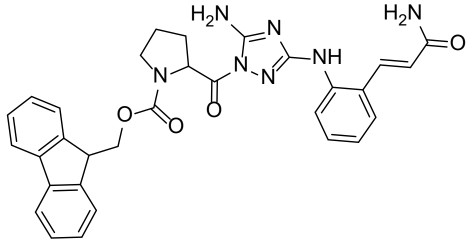

**2710-4**

Chemical Formula: $C_{31}H_{29}N_7O_4$
Molecular Weight: 563.62

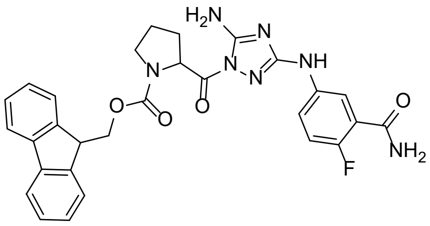

**2710-6**

Chemical Formula: $C_{29}H_{26}FN_7O_4$
Molecular Weight: 555.57

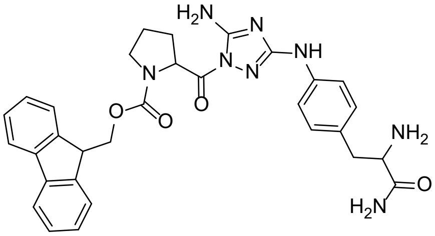

**2710-9,10**

Chemical Formula: $C_{31}H_{32}N_8O_4$
Molecular Weight: 580.65

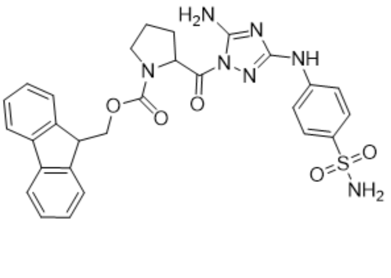

**2710-11,12,13**

Chemical Formula: $C_{28}H_{27}N_7O_5S$
Molecular Weight: 573.63

**2710-14**

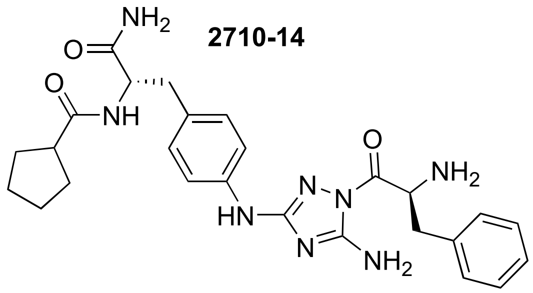

Chemical Formula: $C_{26}H_{32}N_8O_3$
Molecular Weight: 504.60

**2710-15**

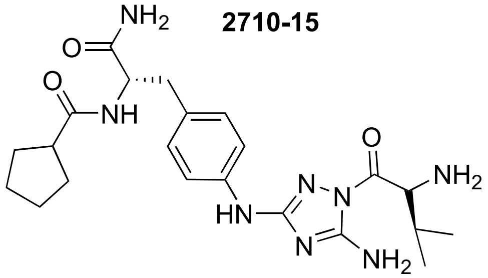

Chemical Formula: $C_{22}H_{32}N_8O_3$
Molecular Weight: 456.55

**2710-16**

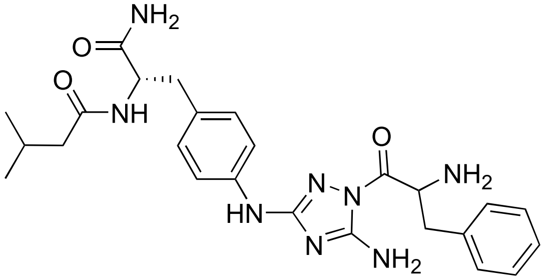

Chemical Formula: $C_{25}H_{32}N_8O_3$
Molecular Weight: 492.58

**2710-17**

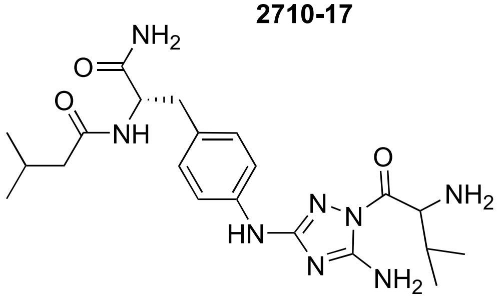

Chemical Formula: $C_{21}H_{32}N_8O_3$
Molecular Weight: 444.54

**2710-18**

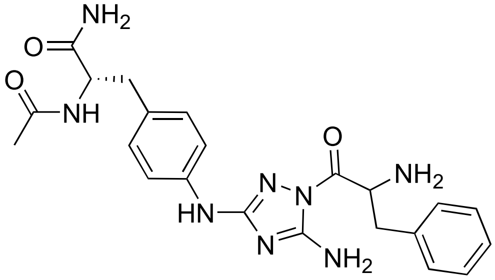

Chemical Formula: $C_{22}H_{26}N_8O_3$
Molecular Weight: 450.50

**2710-19**

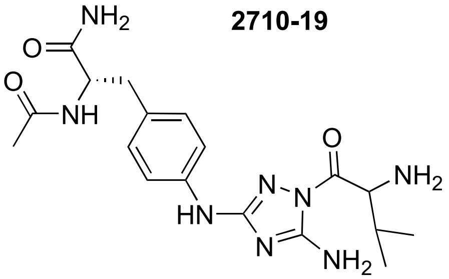

Chemical Formula: $C_{18}H_{26}N_8O_3$
Molecular Weight: 402.46

**2710-20-401**

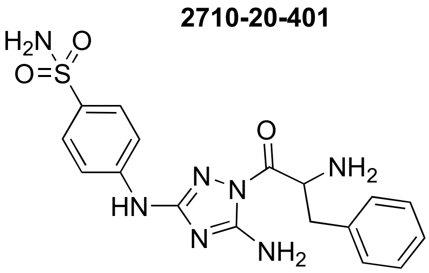

Chemical Formula: $C_{17}H_{19}N_7O_3S$
Molecular Weight: 401.45

**2710-20-526**

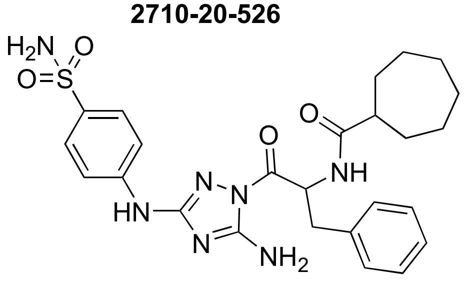

Chemical Formula: $C_{25}H_{31}N_7O_4S$
Molecular Weight: 525.63

**2710-21**

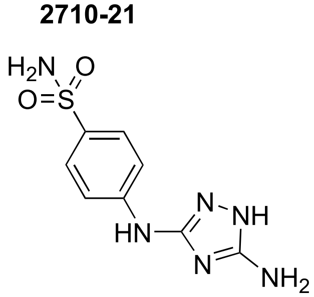

Chemical Formula: $C_8H_{10}N_6O_2S$
Molecular Weight: 254.27

**2710-22**

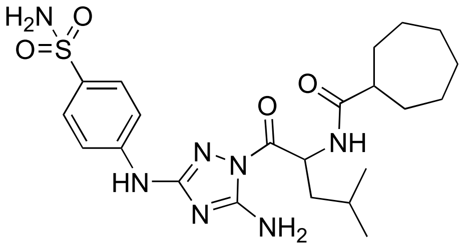

Chemical Formula: $C_{22}H_{33}N_7O_4S$
Molecular Weight: 491.61

**2710-23**

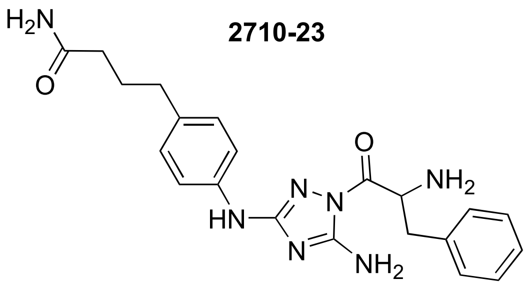

Chemical Formula: $C_{21}H_{25}N_7O_2$
Molecular Weight: 407.48

**2710-24**

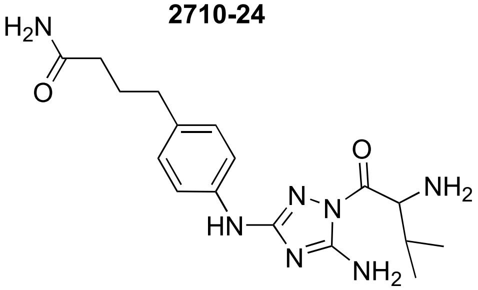

Chemical Formula: $C_{17}H_{25}N_7O_2$
Molecular Weight: 359.43

**2710-25-498**

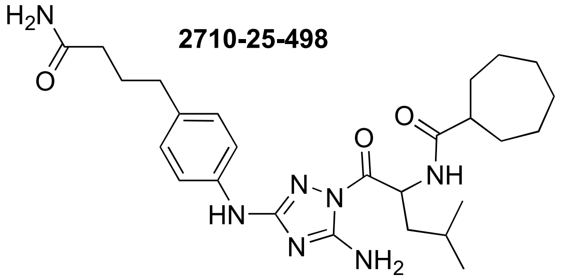

Chemical Formula: $C_{26}H_{39}N_7O_3$
Molecular Weight: 497.64

**2710-25-374**

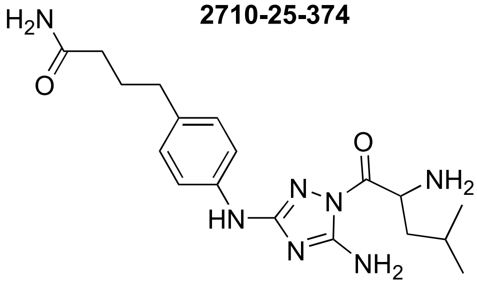

Chemical Formula: $C_{18}H_{27}N_7O_2$
Molecular Weight: 373.46

**27110-26**

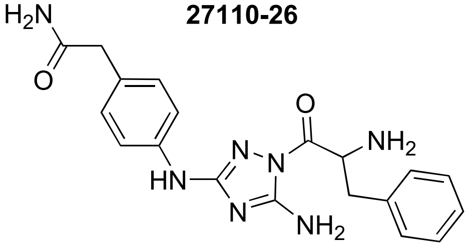

Chemical Formula: $C_{19}H_{21}N_7O_2$
Molecular Weight: 379.42

**2710-27**

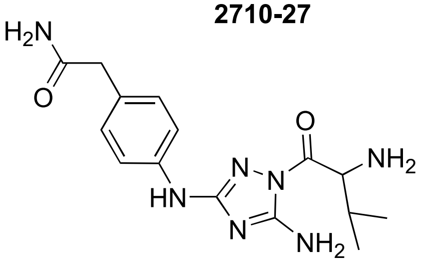

Chemical Formula: $C_{15}H_{21}N_7O_2$
Molecular Weight: 331.38

**2710-28**

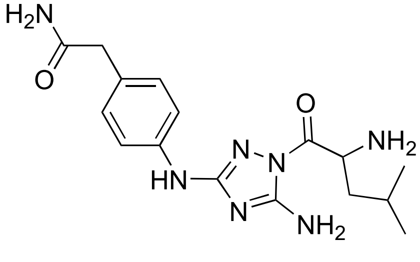

Chemical Formula: $C_{16}H_{23}N_7O_2$
Molecular Weight: 345.41

**2710-30**

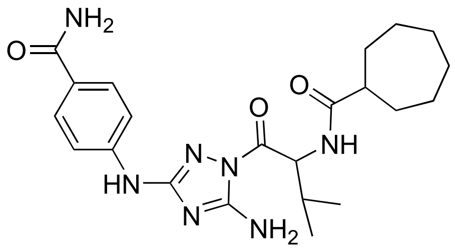


Chemical Formula: $C_{22}H_{31}N_7O_3$
Molecular Weight: 441.54

**2710-31**

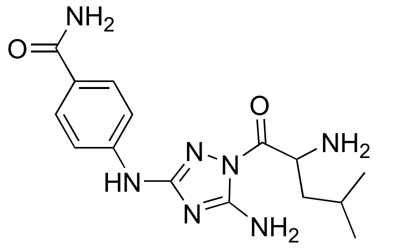


Chemical Formula: $C_{15}H_{21}N_7O_2$
Molecular Weight: 331.38

**2710-32-391**

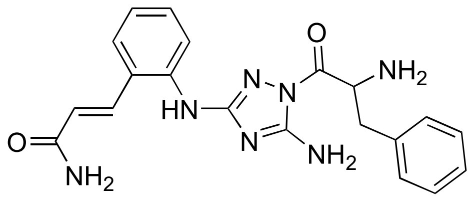


Chemical Formula: $C_{20}H_{21}N_7O_2$
Molecular Weight: 391.44

**2710-32-516**

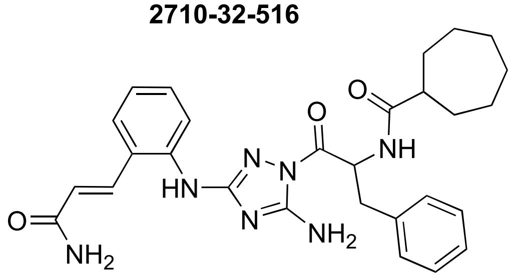


Chemical Formula: $C_{28}H_{33}N_7O_3$
Molecular Weight: 515.62

**2710-33-343**

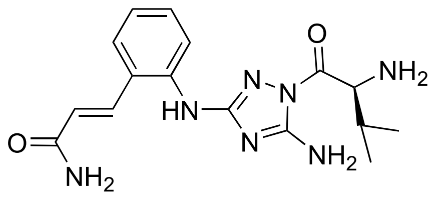


Chemical Formula: $C_{16}H_{21}N_7O_2$
Molecular Weight: 343.39

**2710-33-468**

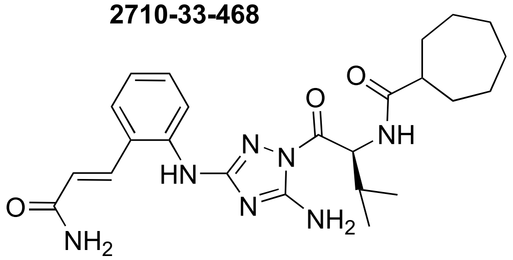


Chemical Formula: $C_{24}H_{33}N_7O_3$
Molecular Weight: 467.57

**2710-34-357**

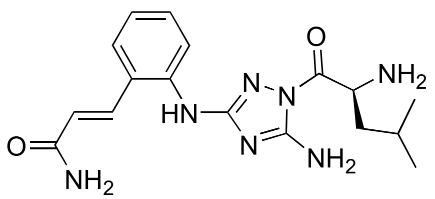


Chemical Formula: $C_{17}H_{23}N_7O_2$
Molecular Weight: 357.42

**2710-34-481**

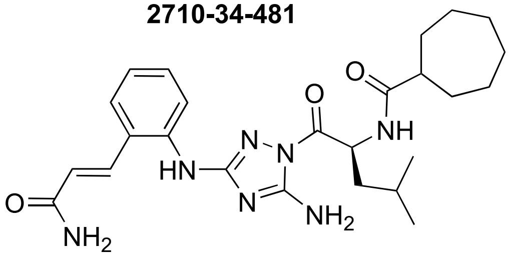


Chemical Formula: $C_{25}H_{35}N_7O_3$
Molecular Weight: 481.60

**Figure S3. Assay of inhibition of human TOP3B relaxation of negatively supercoiled DNA by compounds in the 2710 series.** The compounds were tested at concentration of (a) 200 μM (b) 125 and 62.5 μM. Each panel shows reactions analyzed on the same gel.

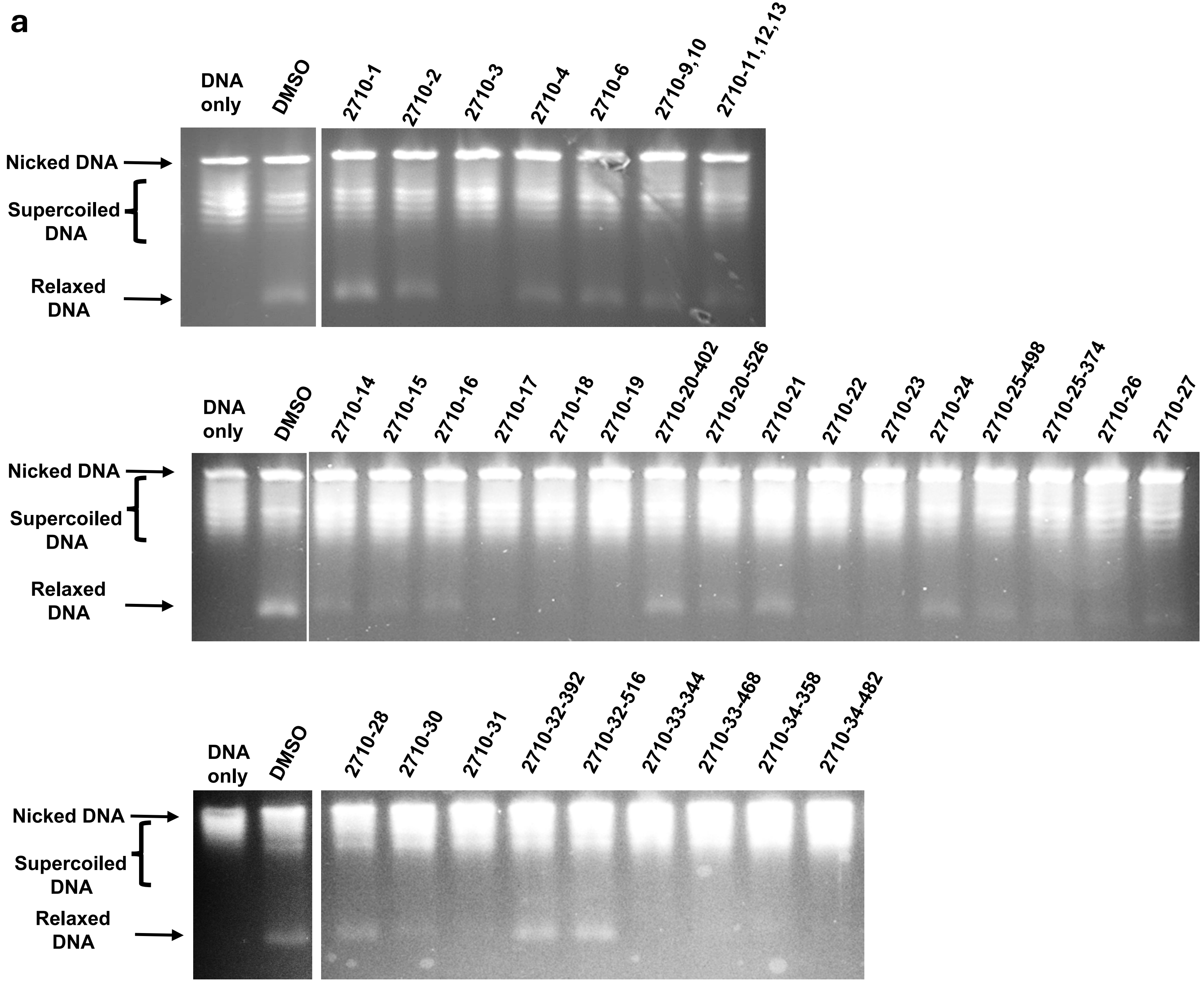

**b**

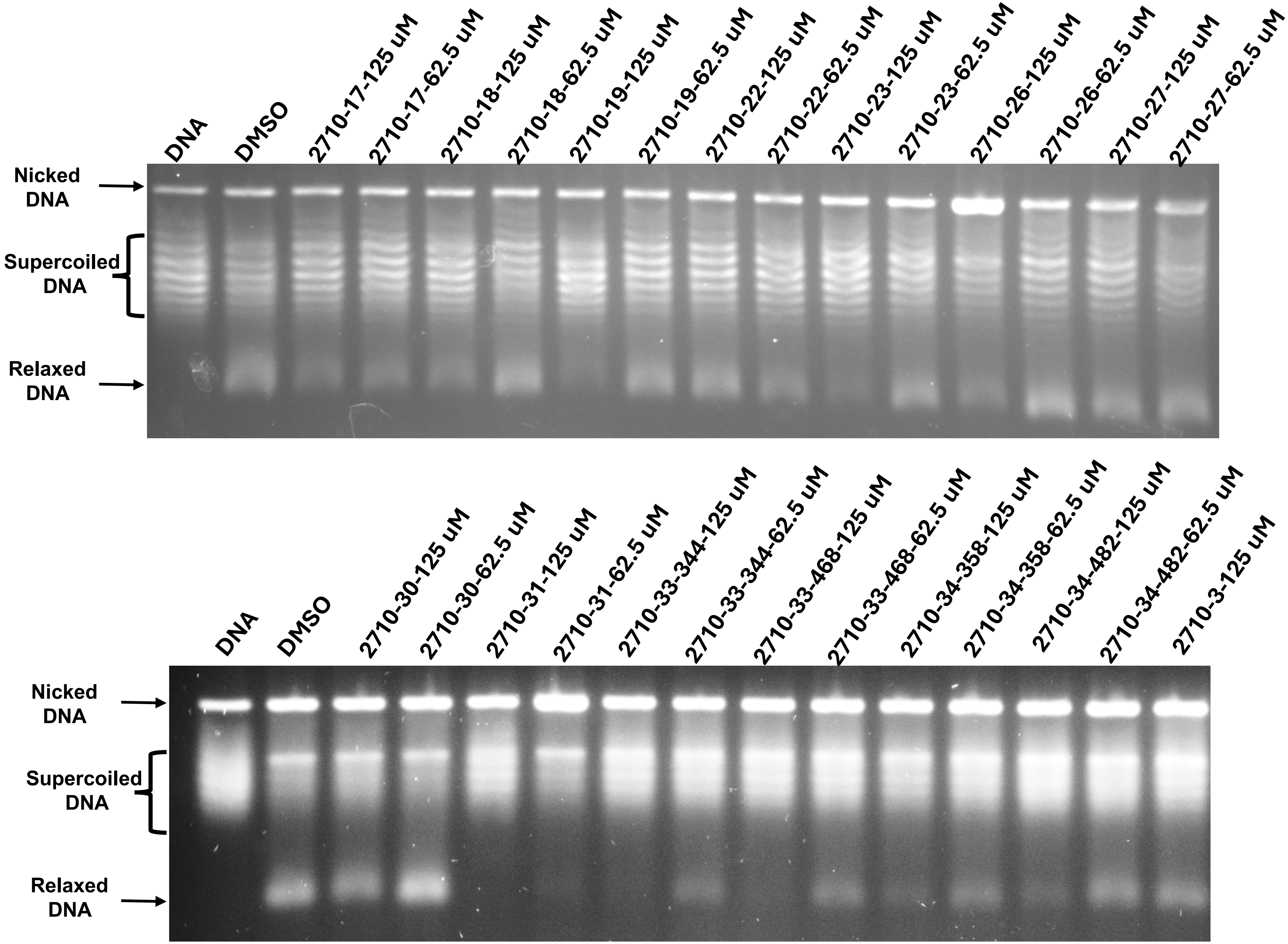
DNA
DMSO
2710-17-125 uM
2710-17-62.5 uM
2710-18-125 uM
2710-18-62.5 uM
2710-19-125 uM
2710-19-62.5 uM
2710-22-125 uM
2710-22-62.5 uM
2710-23-125 uM
2710-23-62.5 uM
2710-26-125 uM
2710-26-62.5 uM
2710-27-125 uM
2710-27-62.5 uM
Nicked DNA
Supercoiled DNA
Relaxed DNA
DNA
DMSO
2710-30-125 uM
2710-30-62.5 uM
2710-31-125 uM
2710-31-62.5 uM
2710-33-344-125 uM
2710-33-344-62.5 uM
2710-33-468-125 uM
2710-33-468-62.5 uM
2710-34-358-125 uM
2710-34-358-62.5 uM
2710-34-482-125 uM
2710-34-482-62.5 uM
2710-3-125 uM
Nicked DNA
Supercoiled DNA
Relaxed DNA

**Figure S4. LC/MS of compound from the 2710 series identified with strongest TOP3B inhibition.**

**2710-31**
**MW331**

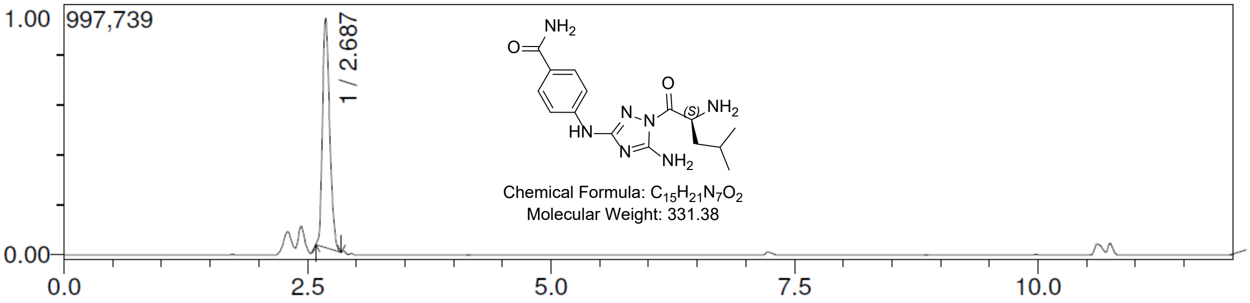


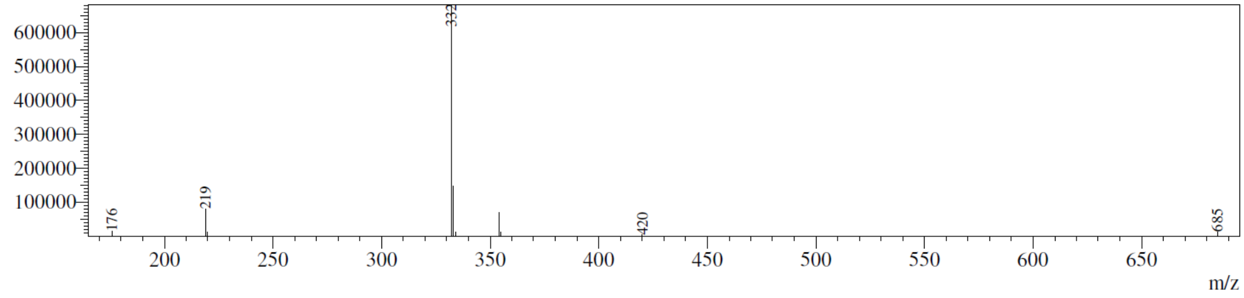


**2710-33-343**
**MW343**

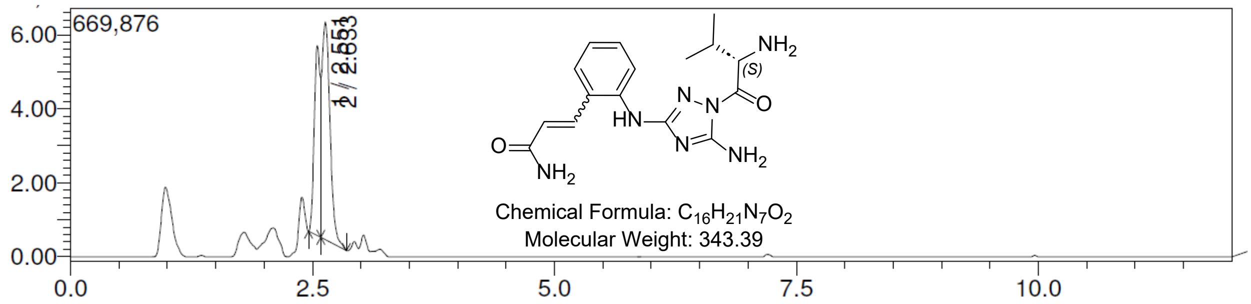


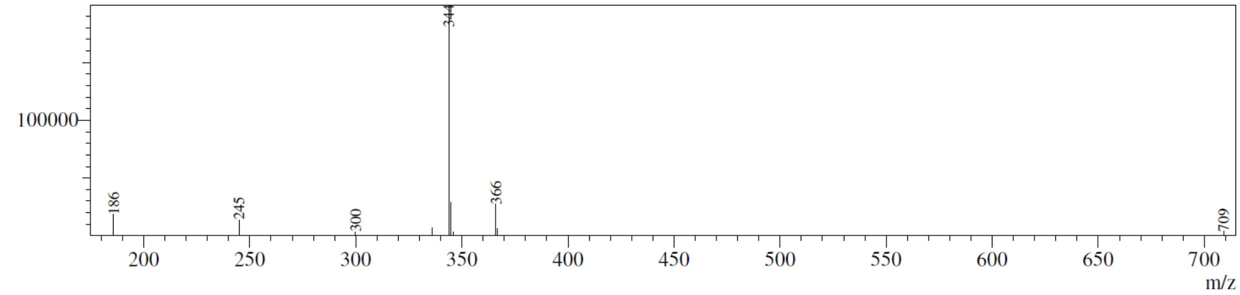


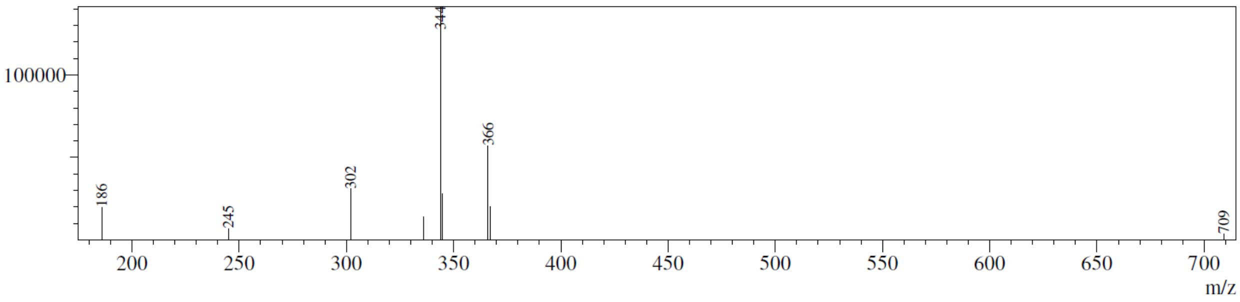

**2710-34-357**
**MW357**

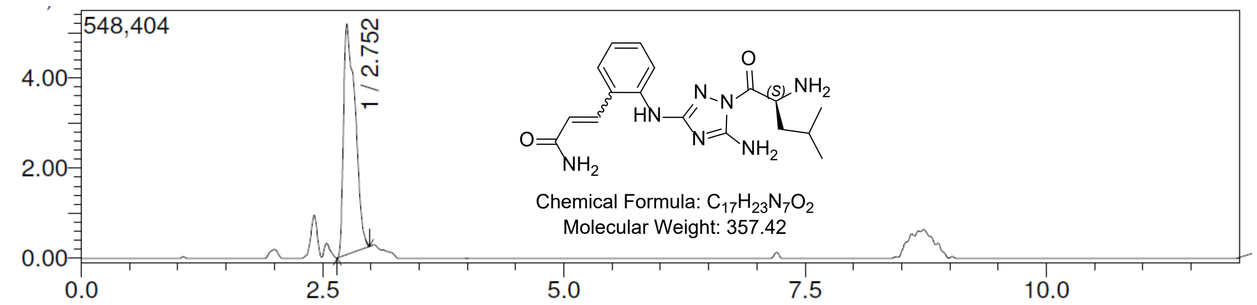


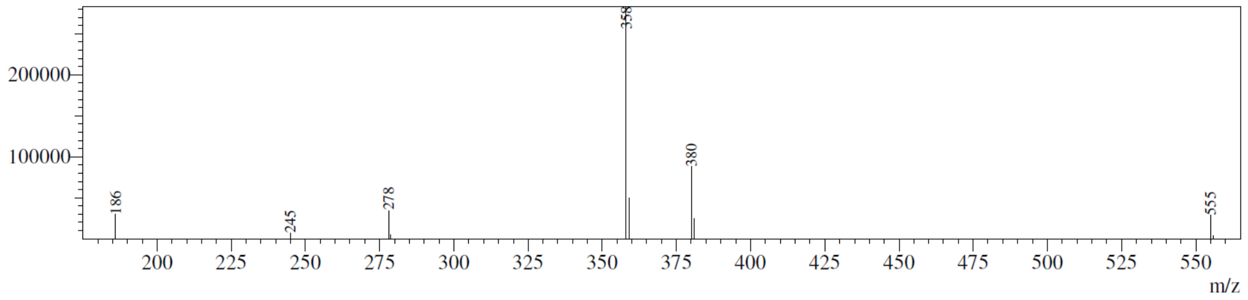


**2710-33-467**
**MW467**

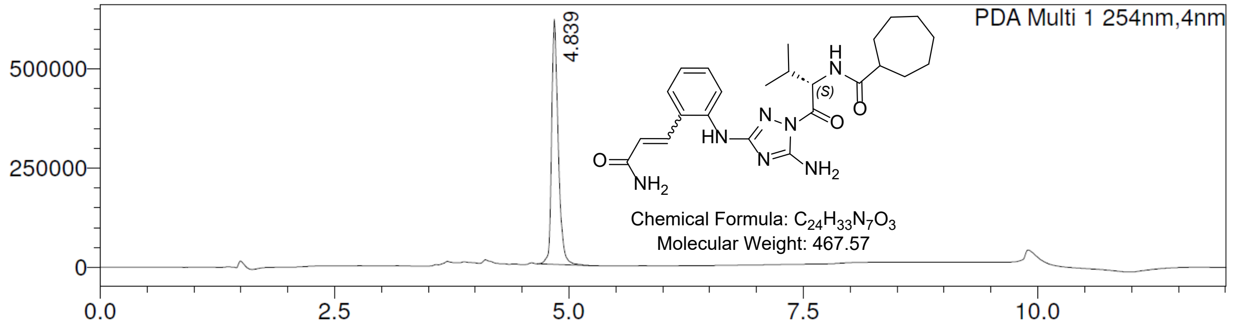


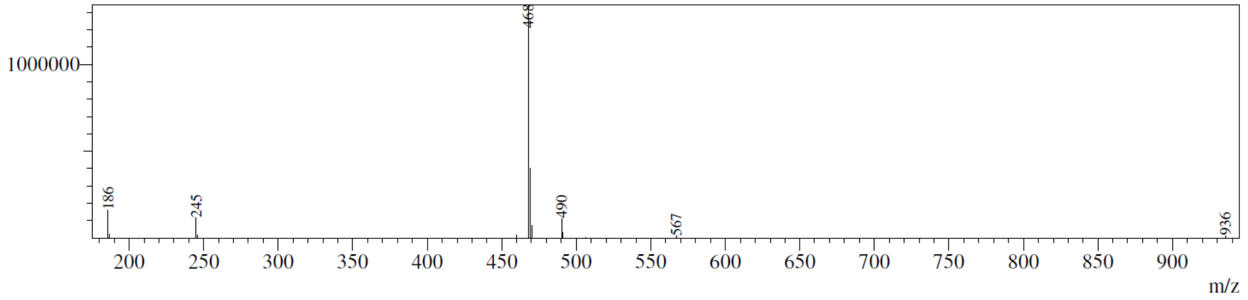

**2710-34-481**
**MW481**

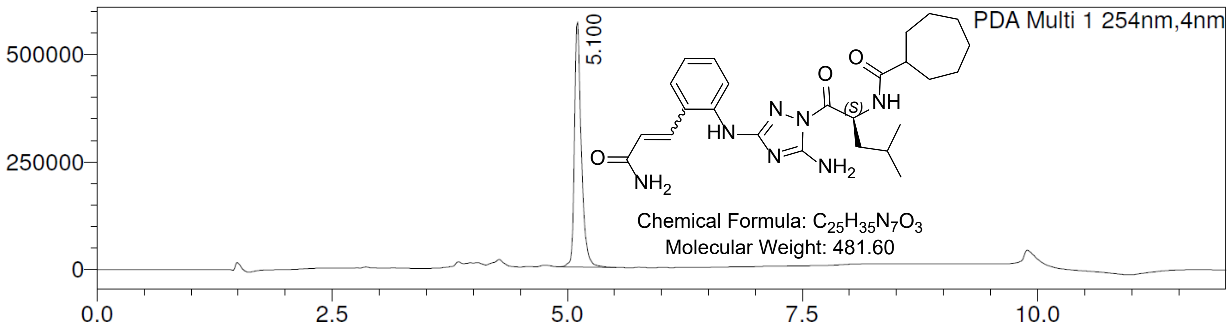


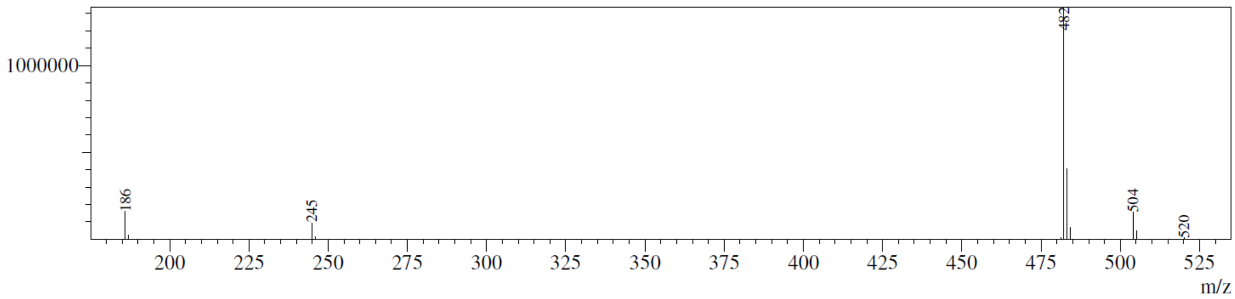
